# When one race is not enough: a relay model explains multisensory response times

**DOI:** 10.1101/2025.07.11.664306

**Authors:** Kalvin Roberts, Thomas U. Otto

## Abstract

Humans typically respond faster to multisensory signals than to their unisensory components, a phenomenon known as the redundant signal effect (RSE). One of the earliest and most influential accounts, the race model, attributes the RSE to statistical facilitation, which arises from parallel, independent processing across sensory modalities. While this model captures some key features of the RSE, it frequently underestimates the observed speed-up leading to violations of the race model inequality (RMI), a benchmark used to test the model’s validity. To reconcile this discrepancy, we introduce the relay model, a minimal extension of the race architecture that incorporates cross-modal initiation. In this model, responses result from two sequential race processes, allowing a signal in one modality to trigger the onset of perceptual decision processing in another. This structure retains statistical facilitation as a core principle while introducing a single free model parameter that divides unisensory processing into gating and decision stages. Through simulations and fits to foundational empirical datasets, we show that the relay model captures both the magnitude and distributional shape of the RSE, including RMI violations. It also accounts for changes in the RSE under asynchronous stimulus onsets, a critical test in multisensory integration research. By extending the classical race model with minimal added complexity, the relay model offers a mechanistically explicit and biologically plausible framework for explaining the dynamics of multisensory decision-making.

**Author Summary:** How does the brain combine information from different senses, like hearing and seeing? For decades, researchers have found that people respond faster when signals from multiple senses occur together than when signals come from a single sense alone. Two main ideas have been proposed to explain this effect: one suggests that the senses operate independently, racing to trigger a response, while the other suggests that they cooperate to enhance processing. In this study, we introduce a new model of multisensory decision-making, called the relay model, which combines elements of both competition and cooperation. We propose that the brain uses a two-stage process: first, the faster sense initiates a second decision process. Then, in that second stage, the faster signal triggers a behavioural response. This relay-like mechanism accelerates responses through competition, while still allowing both senses to cooperate. By capturing key findings from classic experiments, the model offers a new perspective on how the brain combines multisensory information to guide behaviour.

## Introduction

Perceptual decisions are rarely based on information from a single sense. Instead, the brain continuously uses inputs from multiple modalities to guide rapid and adaptive behaviour. This multisensory integration enhances accuracy and accelerates response times (RTs). For instance, if a person both sees an oncoming bicycle and hears its bell, they are likely to react more quickly than if only one of those signals were available. Understanding the computational mechanisms underlying multisensory integration is essential for building models that capture how the brain makes fast and reliable decisions in dynamic environments.

One of the most widely used paradigms to study the benefits of multisensory processing is the redundant signals paradigm [1–4]. In this task, participants are instructed to respond with the same action to different target signals, such as an auditory cue, a visual cue, or both. The critical feature is that signals in the combined condition are redundant, either one alone is sufficient for a correct response. Thus, the task effectively implements a logical OR operation [5–7]. The key finding is that responses to redundant signals are, on average, faster than responses to single signals, a phenomenon known as the redundant signals effect (RSE). The effect has been replicated across a wide range of multisensory pairings (e.g., [8–14]), within single modalities (e.g., [15–19]), across populations (e.g., [20–24]), and under varying task instructions (e.g., [25–28]). This robustness makes the RSE a valuable tool for probing the computational principles of multisensory decision-making.

To understand the mechanisms behind this speed-up, it is useful to first consider how unisensory decisions are modelled. Computational and neurophysiological studies suggest that a key component is the accumulation of sensory evidence over time [29–31]. Specialised neurons modulate their firing in response to stimuli, but because of intrinsic noise, evidence must be integrated over time to improve reliability. Once a decision threshold is reached, a behavioural response is triggered. This process gives rise to typical RT distributions, such as the positively skewed inverse Gaussian (IG) distributions [32, 33], and has proven effective in capturing unisensory decision dynamics [34]. The central question, then, is: how can these models be extended to multisensory decisions?

A milestone answer was offered by Raab [5], who proposed the race model (Fig 1a). Here, two independent evidence accumulation units operate in parallel, each detecting one of the sensory signals. These units are linked by a logical OR gate, meaning that a response is triggered as soon as one or the other unit reaches its decision threshold. When both signals are presented, statistical facilitation occurs: if one process is slower on any given trial, the other may trigger the response first, thereby reducing average RTs (see example trials in Fig 1a). With audio-visual signals (AV), the mechanism of the race model is captured by selecting the smaller of two unisensory RTs (for A and V, respectively):

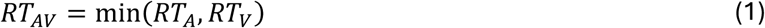

**Fig 1.**
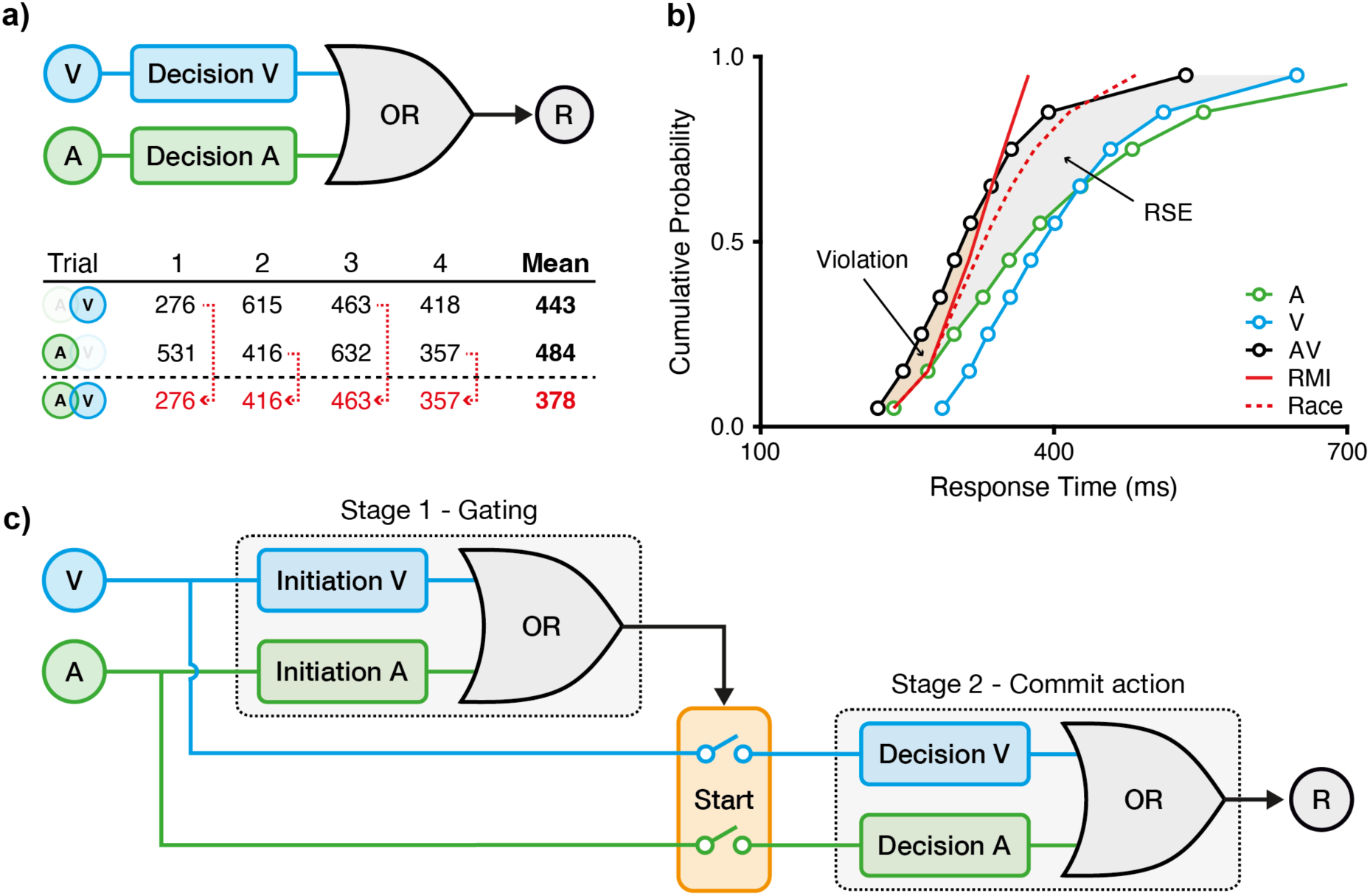
Race models in multisensory decision-making. **a)** Raab’s race model. Auditory (A) and visual (V) signals are processed by two parallel decision units, coupled via a logical OR gate to trigger a response (R). The model predicts faster responses for redundant audiovisual (AV) signals through statistical facilitation, as illustrated by exemplary response times (RTs). On each AV trial, a response is determined by the faster of the two units. Averaging RTs across trials reveals a speed-up of responses for AV compared to unisensory signals. **b)** Miller’s race model inequality (RMI) test. The cumulative RT distribution with redundant AV signals is located to the left of the unisensory RT distributions (A, V), reflecting the redundant signals effect (RSE, grey area). The observed RSE exceeds the prediction of Raab’s race model, including significant violation of the RMI (orange area), indicating a processing interaction between audition and vision. Data replotted from [4]. **c)** Relay model. To reconcile Raab’s race model with RMI violations, we propose a two-stage model comprising two sequential race processes. As an audiovisual processing interaction, the first stage provides a start signal for the second stage, thus functioning as a gating mechanism that allows for cross-modal initiation. The second stage operates analogously to Raab’s race model to initiate a response.

Consequently, the cumulative distribution function (CDF) for RTs in the redundant condition follows probability summation, where the chance of responding by time *t* increases by having two independent chances to respond:

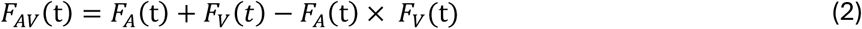

Here, F_AV_(t) is the cumulative probability of responding at time t to both signals, and F_A_(t) and F_V_(t) are the unisensory RT distributions. Thus, race models provide parameter-free predictions of the RSE based on unisensory data, offering concrete and testable predictions about the expected magnitude of the RSE and the conditions under which it should be most pronounced.

Building on probability summation in the race model, two key principles to predict the expected speed-up emerge [35]. First, the *principle of congruent effectiveness* predicts that the RSE is largest when unisensory RT distributions overlap. This overlap can be optimised by introducing a temporal delay between signals, known as stimulus onset asynchrony (SOA), which can offset inherent RT differences. For instance, if auditory responses are on average 30 ms faster, delaying the auditory relative to the visual signal by 30 ms aligns the two unisensory RT distributions and maximises the RSE (e.g., [2, 35, 36]). Second, the *variability rule* states that the RSE is driven by the variability of RTs in unisensory conditions, which can be modulated for example through signal intensity. As weaker signals tend to produce greater RT variability, the RSE is expected, and has been shown, to increase under such conditions (e.g., [35, 37]). Together, these findings show that the race model captures key patterns of multisensory facilitation, reinforcing its value as a simple yet powerful explanatory framework.

Despite their predictive power, race models have clear limitations. As a second milestone, Miller [4] demonstrated that the RSE often exceeds what can be explained by probability summation alone. Allowing for a maximally negative correlation between unisensory processes [38], the probability of responding to redundant signals cannot exceed the sum of the individual unisensory probabilities:

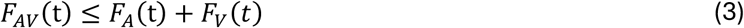

This so-called race model inequality (RMI) provides a testable boundary for race model predictions (Fig 1b). Empirical RMI violations demonstrate that the observed RSE cannot be attributed to the race model alone, implying some interaction between sensory channels. Given that RMI violations have been reported consistently, it has contributed to a widespread rejection of the race model architecture in explanations of the RSE (for exceptions, see [11, 39]).

Importantly, the RMI, and race models in general, implicitly assume a single-stage architecture: two isolated, parallel accumulation processes leading to the same response (for comprehensive discussions of the underlying assumptions, see [7, 40, 41]). This assumption is analytically convenient but biologically simplistic. As soon as the decision process includes sequential stages with crossmodal interactions, the foundational assumptions of the RMI framework begin to break down [42]. This raises a critical question: how can multisensory decision-making models be extended to capture biologically plausible multi-stage dynamics?

In unisensory decision-making, multi-stage models are increasingly prevalent. For example, Carpenter et al. [43] proposed a two-stage architecture in which early detection units trigger a later accumulation unit (see [44–46]). Other work suggests that evidence accumulation is itself gated by an initial selection process (e.g., [47–50]). In these multi-stage models, the total RT is expressed as the sum of processing times associated to individual stages (e.g., RT_A_ = T_A1_ + T_A2_ for an auditory stimulus). While such multi-stage frameworks are well established in unisensory contexts, their extension to multisensory decision-making remains comparatively underexplored (but see [51]).

In this study, we develop and test a relay model to address this gap in multisensory decision making (Fig 1c). This model architecture features two sequential race units, allowing a signal from one modality to initiate evidence accumulation in the other. For instance, a fast visual initiation signal (*T_V_*_1_) might trigger auditory evidence accumulation (*T_A_*_2_), and vice versa. As a result, the total multisensory RT is given by the sum of two race processes:

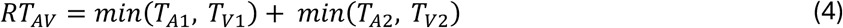

Crucially, we show in the following that RMI violations, previously unexplained by the race model, emerge naturally from the relay architecture. Furthermore, the model not only accounts for the scale of the RSE but also captures dynamics when the timing between signals is manipulated. Thus, by reconciling the milestone contributions by Raab and Miller, it offers a biologically plausible, computationally grounded framework that reflects the staged nature of perceptual decisions in a multisensory context.

## Results

### The relay model explains multisensory processing dynamics

To evaluate the plausibility of the relay model, we tested whether its predictions align with empirical RTs reported in Miller’s benchmark dataset (Fig 1b in [4]). As a key advancement for the understanding of multisensory processing, the model incorporates that perceptual decision processes comprise sequential processing stages. For example, an auditory RT is modelled as the sum of two stages (e.g., RT_A_ = T_A1_ + T_A2_).

To quantify the contribution of each stage, we introduce the RT share parameter, which expresses the duration of each stage as a proportion of the total RT. For instance, if RT_A_ = 400 ms and the first stage (T_A1_) lasts 80 ms, then T_A1_ contributes 20% of the total RT (Fig 2a, left). We formalised the model by fitting IG distributions to the unisensory RT data. This distribution was selected for two reasons. First, its positive skew and non-negative support closely match the empirical RT distributions observed in simple detection tasks [32, 33]. Second, under specific conditions, the sum of two IG-distributed variables remains IG-distributed, allowing the total RT to be modelled as the sum of two stages while maintaining analytical tractability. The RT share thus expresses the relative contribution of two sequential IG-modelled stages.

**Fig 2.**
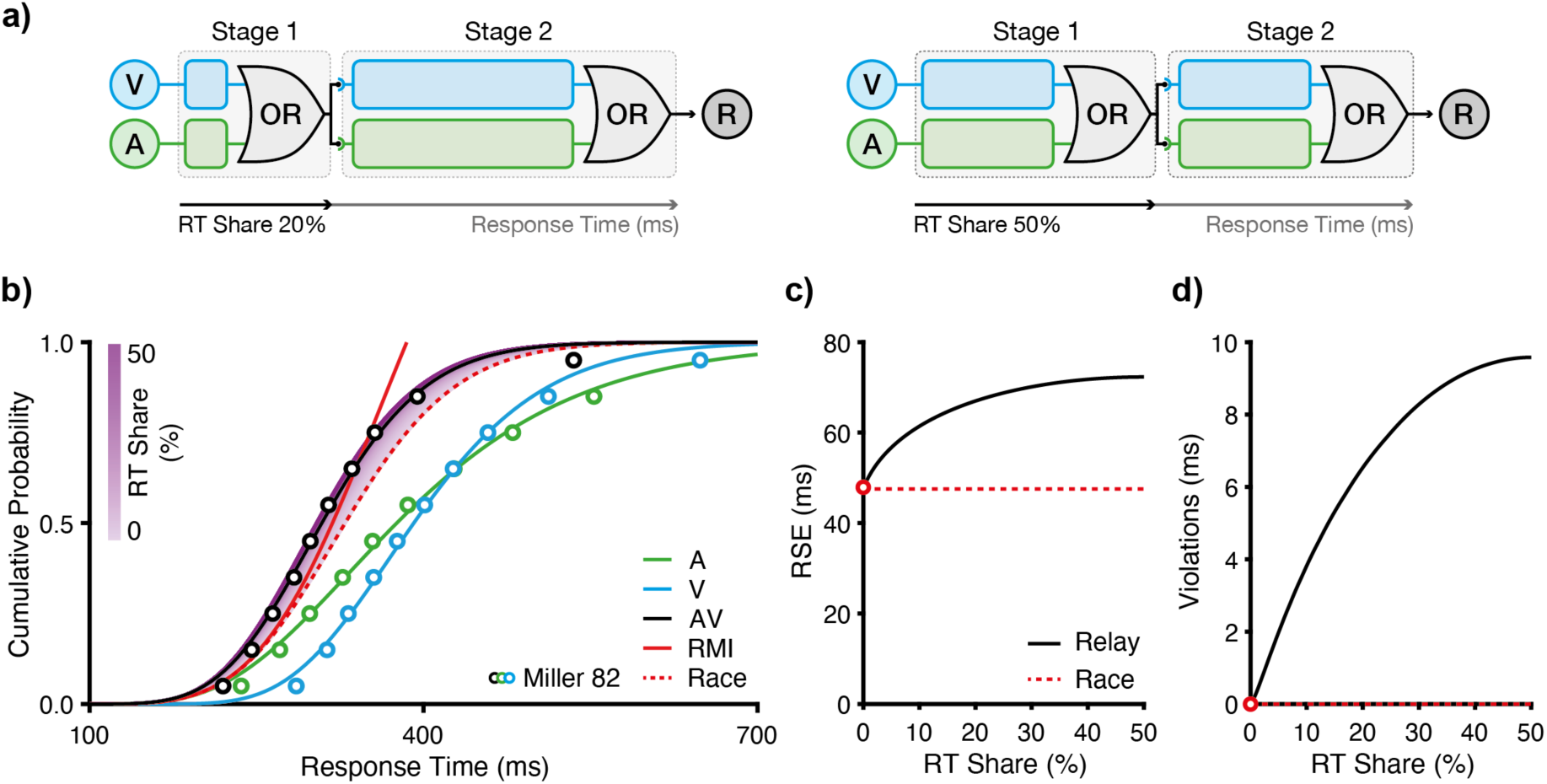
Predictions of the relay model. **a)** RT share. In the relay model, the RT share denotes the proportion (%) of the overall RT attributed to the first processing stage: 20% (left) and 50% (right). Schematic diagrams correspond to the model architecture introduced in Fig 1c. **b)** Modelling based on data from Miller [4]. The relay model is constrained by the RT distribution for auditory (A) and visual (V) signals (Table 1). The shaded region depicts relay model predictions for RT shares ranging from 0% to 50%. The empirical RT distribution for redundant audiovisual (AV) signals is best fit with an RT share of 22.6%. Circles represent the same data shown in Fig 1b. **c)** RSE and **d)** RMI violations as a function of RT share. The relay model predicts RSEs that exceed those of Raab’s race model, including systematic RMI violations. Predictions decline symmetrically for RT shares above 50% (e.g., RT shares of 20% and 80% yield identical predictions).

**Table 1.** Parameters for the best-fitting unisensory IG distributions.

| Data | Auditory |  | Visual |  |
| --- | --- | --- | --- | --- |
| | $\mu_A$ (ms) | $\lambda_A$ (ms) | $\mu_V$ (ms) | $\lambda_V$ (ms) |
| Miller (1982) | 398.6 | 3272.4 | 401.9 | 7308.6 |
| Miller (1986) | 231.0 | 3930.6 | 348.0 | 4979.2 |

To constrain the model, we fitted IG distributions to unisensory auditory and visual RTs reported in Figure 1 from [4]. We estimated parameters by minimising the root mean squared error (RMSE) between the empirical and model CDFs (Methods; Table 1). At this stage, no decomposition into stages is required; the total RT is modelled directly.

We then extended the model to the multisensory case. The relay model assumes a non-competitive relay architecture (Fig 1c; Eq 4). During the first stage, unisensory processes (e.g., T_A1_, T_V1_) race to completion. Passing on the baton, the winner then initiates the next stage, triggering a race between the second-stage components (e.g., T_A2_, T_V2_). The winner of this second race determines the motor response.

In this framework, the RT share governs the relative durations of the two stages, shaping how the system transitions across sensory modalities and stages. Crucially, this architecture enables cross-modal initiation: for example, a visual stage 1 winner can trigger an auditory stage 2 process. This feature distinguishes the relay model from classical race models. For simplicity, we used a single RT share parameter for both modalities; however, the model can be extended to support distinct RT shares per modality, reflecting differences in sensory latencies.

To examine how the RT share affects model behaviour, we simulated the relay model across RT share values for stage 1 from 0% to 50% (Fig 2b). As the two stages add up to 100%, this range encompasses the whole dynamic space (values greater than 50% are symmetric). At 0% RT share, the relay model reduces to the classical race model (Fig 1a; Eq 1) which predicts an RSE of 48 ms. In contrast, relay model predictions amplify the RSE up to 72 ms (1.5 * the race model) at 50% RT share (Fig 2c). The race model also predicts no violations of the RMI, but with an RT share of 50%, the relay model predicts RMI violations in the order of 10 ms (Fig 2d). Thus, the RT share, as an interpretable single-parameter implementation of the dual-stage architecture, enables the relay model to capture the key multisensory effects observed in human behaviour.

To evaluate the model’s fit to empirical data, we estimated the optimal RT share by minimising the RMSE between empirical and predicted multisensory CDFs. For parsimony, we again constrained auditory and visual RT shares to be equal. The optimal RT share was 22.6% (RMSE = 0.023), which corresponds to first-stage durations of 89.87 ± 31.37 ms (Mean ± SD) for audition and 90.62 ± 21.25 ms for vision. The model captures the distributional shape of the RSE virtually perfectly.

In summary, with only one additional degree of freedom beyond the unisensory fits, the relay model successfully reproduces two hallmark signatures of multisensory integration: larger RSEs than predicted by the race model, including RMI violations. These findings demonstrate that a simple relay architecture offers a parsimonious yet powerful framework for modelling the dynamics of multisensory processing.

### Predictions of multi-stage relay models

While our initial analyses focused on a two-stage relay architecture, the model is not inherently restricted to two processing stages. In principle, multisensory decision-making could involve three or more stages, with each stage comprising a race between modalities (Fig 3a). This raises a key question: how many relay stages account for empirical multisensory RTs best?

**Fig 3.**
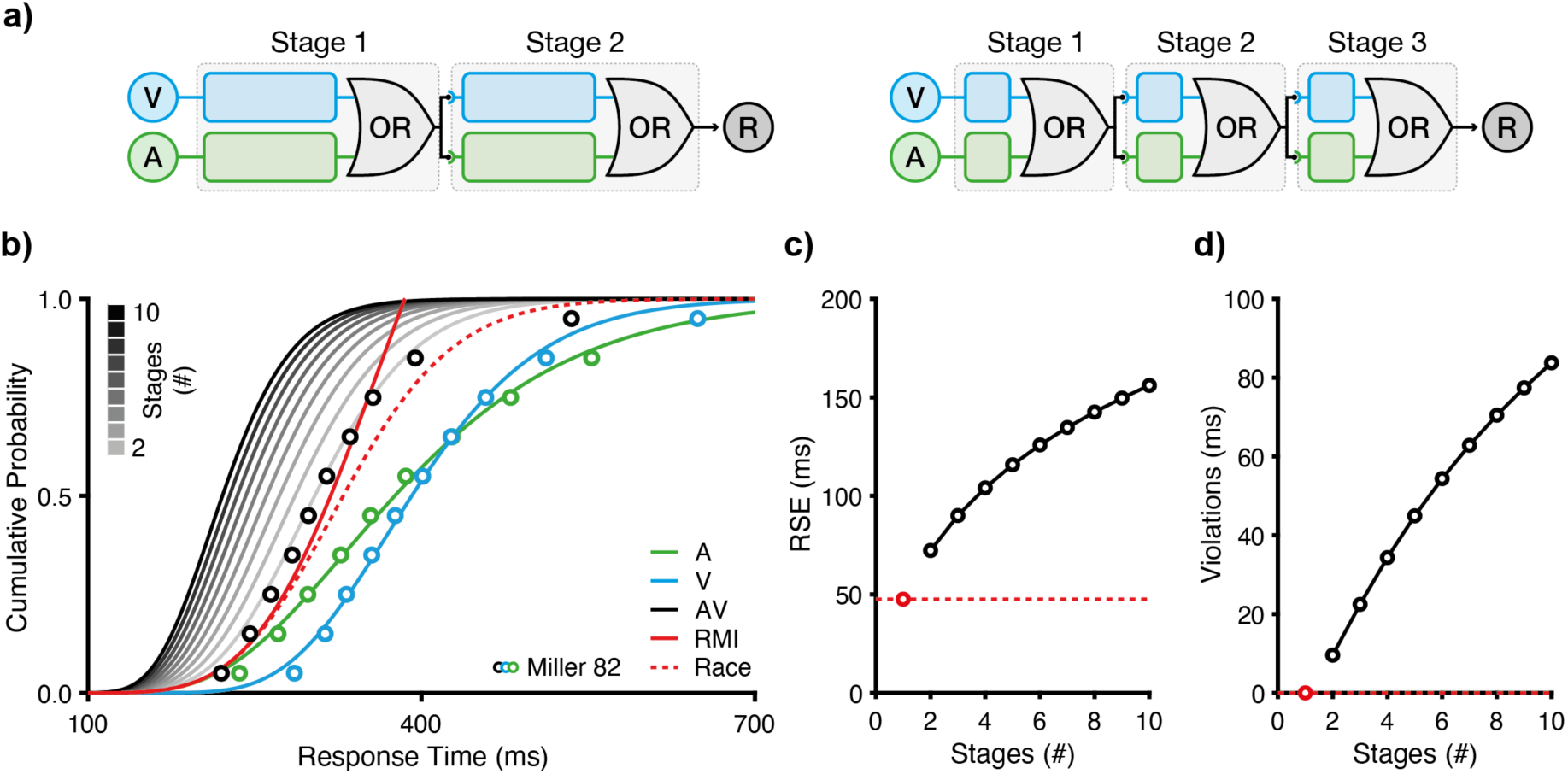
Multi-stage relay models. **a)** Number of stages. Relay models with two and three sequential stages are shown. Each stage passes a start signal to the next, with the overall RT evenly distributed across stages. The two-stage schematic diagram corresponds to the architecture introduced in Fig 1c. **b)** Modelling based on data from Miller [4]. Greyscale lines depict predicted cumulative RT distributions for increasing numbers of stages, from two up to ten. Circles represent the same data shown in Fig 1b. **c)** RSE and **d)** RMI violations as a function of the number of relay stages. Relay models with three or more stages predict RSEs and RMI violations that far exceed empirical observations. Predictions from Raab’s race model (one stage) are indicated by red circles for reference.

To address this, we simulated relay architectures with one to ten stages, using the same unisensory IG fits to constrain each model (Fig 3b). In every simulation, the total RT was evenly divided across N stages, such that each stage was allocated 1/N of the total RT. Accordingly, the RT share was held fixed at this proportion (i.e., not optimised). This design enabled us to assess how increasing the stage count affects predicted RSEs and RMI violations, while holding model complexity and parameterisation constant.

Simulation results show that both predicted RSEs and RMI violations increase with stage count, but with diminishing returns (Fig 3c,d). The most substantial gain occurs between the race model and the two-stage relay model. Adding more stages yielded progressively smaller gains. Crucially, higher stage counts led to increasingly exaggerated predictions. For example, the ten-stage model predicted an RSE of 156 ms and RMI violations of 84 ms, which far exceed empirically observed values.

To evaluate these architectures quantitatively, we compared predicted to empirical multisensory CDFs (Fig 3b). Because the RT share was fixed (rather than optimised) and all models shared identical unisensory parameters, this provided a fair, complexity-controlled comparison. The two-stage relay model emerged as the best configuration, yielding the lowest RMSE (RMSE = 0.030), more than twice as accurate as the one-stage race model (RMSE = 0.085).

Adding further stages degraded model performance (e.g., three-stage RMSE = 0.102; ten-stage RMSE = 0.399), consistent with their overestimation of RSE and RMI effects.

In summary, while relay architectures can, in principle, support an arbitrary number of stages, the two-stage configuration strikes the optimal balance. It captures observed multisensory enhancements without inflating predictions beyond empirical observations.

### Relay model and the temporal dynamics of multisensory decisions

Having established that the two-stage relay model captures both RSE and RMI violations under standard conditions, we next examined its ability to capture multisensory behaviour under conditions of temporal asynchrony. Specifically, we simulated SOA conditions, in which one sensory signal leads the other (Fig 4a). As a hallmark manipulation in multisensory research, SOA manipulations provide a stringent test of multisensory models by probing how response dynamics adapt when stimuli are no longer presented simultaneously [e.g., 36, 52, 53].

**Fig 4.**
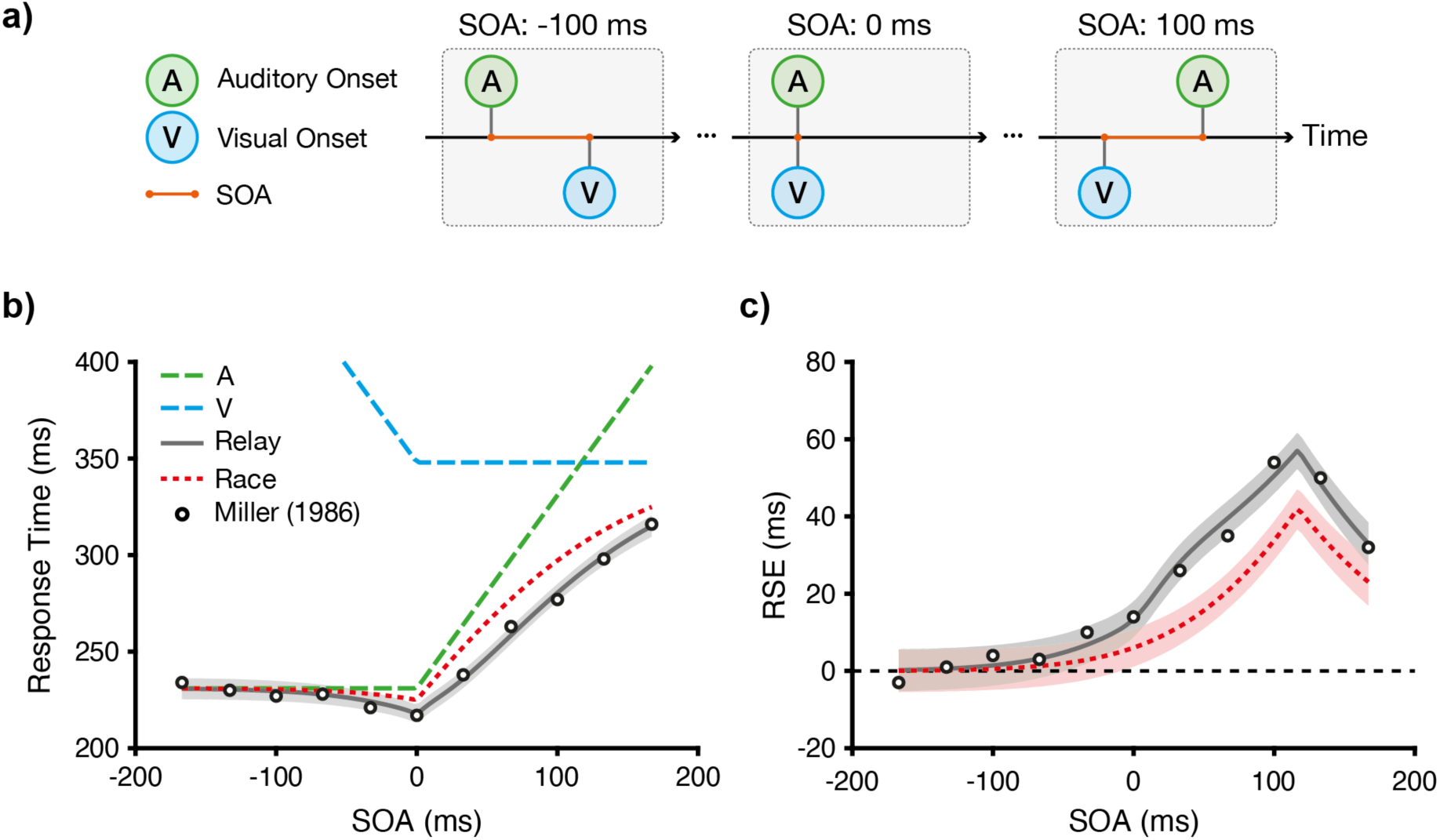
Temporal dynamics in multisensory processing. **a)** Stimulus onset synchrony (SOA). *Left:* negative SOAs represent visual lags relative to auditory signal onset. *Centre:* an SOA of 0 ms represents synchrony between the auditory and visual onsets. *Right:* positive SOAs represent auditory lags relative to visual onset. **b)** Modelling based on data from Miller [36]. Dashed lines represent unisensory RTs as reference, adjusted for SOA (e.g., the auditory delay is added to auditory RTs for positive SOAs). Circles show empirical RTs to redundant AV signals across SOAs, which are faster than predictions from Raab’s race model (red dotted line). The relay model fits the empirical RTs with an RT share of 13.5%. **c)** Modulation of the RSE across SOAs. Red and grey lines show predictions from Raab’s race model and the relay model, respectively. Both models predict the maximal RSE around SOAs of ~100 ms, but empirical RSEs consistently exceed race model predictions. Shaded areas show 95% confidence intervals derived from bootstrap procedures.

We used the benchmark dataset from Miller [36], which systematically varied SOAs between auditory and visual signals from −167 ms (visual lead) to +167 ms (auditory lead) in 33 ms increments. This dataset exhibits two characteristic features. First, the fastest multisensory responses occur under synchronous conditions (SOA = 0 ms; Fig 4b), as any deviation from synchrony introduces delay. Second, the largest RSE is observed when the SOA compensates for the difference in unisensory RTs, a phenomenon introduced earlier as the principle of congruent effectiveness [35]. In Miller’s data, visual RTs were ~100 ms slower than auditory RTs. Accordingly, the largest RSE was observed when the auditory signal lagged by ~100 ms (Fig 4c). While the classic race model captures these characteristics, it underestimates RSE magnitudes across SOA conditions.

To accommodate temporal asynchrony in the relay model, we implemented a simple extension. Because a signal cannot be processed before it is presented, we added a delay to the corresponding processing stage. Thus, stage 1 processing for each modality begins with signal onset. For example, if the auditory signal is delayed by Δ*t*, the stage 1 race completion time becomes *T*_1_ = *min*(*T_A_*_1_ + Δ_t_, *T_V_*_1_). Critically, we retained the relay model’s core feature of cross-modal initiation: stage 2 processing is triggered by the first modality to complete stage 1 (Fig 1c). If stage 2 is initiated before the delayed signal is physically present, processing of that modality is further delayed until its onset. In the model, we simply added the residual delay at stage 2. This mechanism becomes increasingly important at larger SOAs, where the lagging modality may be partially or fully excluded. When the SOA exceeds the combined duration of both stages, the model response effectively reduces to the unisensory condition of the leading signal.

We tested whether this simple model extension could fully reproduce the empirical data. First, we constrained the model by fitting its unisensory parameters (Methods; Table 1). With these parameters fixed, we optimised the model’s single free parameter, the RT share, by minimising the RMSE between predicted and observed RSEs across SOA levels. The best-fitting RT share was 13.5% (RMSE = 2.5 ms), corresponding to first-stage durations of 31.06 ± 7.53 ms for auditory and 46.79 ± 12.37 ms for vision, and produced an excellent fit with all data points within 95% confidence intervals.

Thus, the relay model successfully captured both qualitative and quantitative features of the data. It predicted the fastest RTs at synchronous onset, identified the SOA that maximised the RSE, and closely matched the RSE magnitudes across all SOA conditions (Fig 4b,c). Thus, by incorporating stage-based architecture that allows for cross-modal initiation, the relay model offers a mechanistically grounded account of how temporal asynchrony shapes multisensory decision-making.

## Discussion

The robust speed-up of responses observed for multisensory compared to unisensory signals provides a critical window into the computational strategies underlying perceptual decision-making. The RSE not only reflects a prominent performance advantage but also constrains the set of plausible models that explain how the brain integrates information across modalities. As one of the earliest and most influential frameworks, the race model [5] proposes that responses are initiated by whichever unisensory channel first reaches a decision threshold, assuming that modalities operate independently in parallel (Fig 1a). While this model largely accounts for the RSE through statistical facilitation, it typically underestimates the observed behavioural enhancement. Most notably, it imposes the RMI as an upper bound on the RSE, which is frequently violated under multisensory conditions (as demonstrated in numerous follow-ups to [4]; Fig 1b). These systematic violations strongly argue against purely parallel architectures and have led many to favour explanations in which multisensory inputs are integrated into a single decision process (e.g., [54–57]).

To reconcile the conceptual parsimony of the race model with empirical RMI violations, we introduced the relay model, a minimal extension that includes two race-like stages (Fig 1c). As a key feature, the relay model allows for cross-modal initiation in addition to statistical facilitation. In its most basic form, the model includes a single free parameter, the RT-share, which partitions the overall RT between the processing stages (in terms of both mean and variance). Our simulations and fits to classical data [4] demonstrate that the relay model captures the magnitude of the RSE, including RMI violations, while closely approximating the observed RT distribution (Fig 2), with a two-stage architecture being fully sufficient (Fig 3). Moreover, simulations and fits to data [36] show that it captures the RSE modulation under asynchronous signal presentation (Fig 4). Together, these findings indicate that a two-stage relay configuration is both computationally parsimonious and descriptively powerful, providing an efficient and accurate framework for modelling multisensory decision-making.

The relay model does not commit to any specific cognitive mechanism underlying the two stages. One speculation we have presented here is the idea that modality-specific signals race to trigger a gate that, once opened, enables evidence accumulation processes to begin. The concept of gated accumulation is well-established in perceptual decision-making. For example, gated accumulation mechanisms have been applied to unisensory decision processes [e.g., 49, 58, 59], where gating determines the onset of evidence accumulation and selects which sensory signals contribute. Crucially, this framework is supported by physiological evidence: early event-related potential (ERP) components have been shown to predict the initiation of accumulation [48, 60]. In the relay model, a fast visual initial stage may open the gate for the accumulation of auditory evidence, a process that would be beneficial when the signals are expected to co-occur. In the context of multisensory ERP studies, multisensory modulation of ERPs has been observed as early as 40 to 80 ms after stimulus onset [61–63]. These latencies correspond well with our predicted first-stage components, ranging from 30 to 90 ms across both fitted datasets [4, 36]. Interestingly, some evidence suggests that early multisensory integration effects predominantly enhance neural activity in the nondominant sensory modality [62]. Given auditory dominance (i.e., faster and more accurate), the relay model would predict that the auditory process wins the first race more often. Given this imbalance, the visual second stage would benefit most from cross-modal initiation and would likely lead to the observed enhancement in the non-dominant modality. Of course, for visual dominance, the opposite will be true. These ERP markers, therefore, provide plausible neural correlates for the first model stage, thereby linking the relay architecture to measurable brain dynamics. Future work could test this interpretation more directly through a joint modelling of behaviour and neural recordings, potentially disentangling the first and second phases temporally and across sensory modalities.

In the context of multisensory modelling, the time-window-of-integration (TWIN) model introduced a two-stage architecture comprising an initial sensory processing stage followed by a pooling stage that integrates signals to accelerate responses under multisensory conditions [27, 51]. The TWIN model accounts for intersensory facilitation effects with its initial stage, for example, when an auditory distractor facilitates responses to a visual target despite being task-irrelevant. Similarly, Nickerson [64] attributed such effects to enhanced response preparedness whereby, for example, a fast auditory signal primes the system to respond more rapidly to a coincident visual stimulus. These frameworks provide important conceptual precedents for separating stages in models of multisensory decision-making, motivating the architectural structure adopted by the relay model.

The first stage of the relay model, in fact, closely aligns with the TWIN model, implementing a mechanism we refer to as cross-modal initiation: the signal that completes the initial gating stage first triggers the onset of processing in the subsequent main decision stage. The key innovation of the relay model lies in formalising both stages within a race-like architecture, which preserves statistical facilitation, as originally proposed by Raab [5], as the central mechanism driving the RSE. By integrating concepts of intersensory facilitation and race models, the relay model offers a mechanistically explicit and testable account of multisensory enhancements, including RMI violations.

One of the strengths of Miller’s work is that the RMI provides a distribution-free test of Raab’s basic race architecture directly using empirical RT data. While our implementation of the relay model uses IG distributions for simulation and fitting, we emphasise that the model’s core properties, and in particular its ability to violate the RMI, do not depend on this specific distributional choice. In fact, the key mechanism underlying RMI violations in the relay model is cross-modal initiation, where the second decision stage is triggered by whichever modality completes the initial stage first. This feature can be demonstrated using a variety of distributional choices, underscoring the generality of the model’s architecture.

One limitation of the current relay model is that it does not account for trial history effects, which have been shown to influence RTs in RSE experiments [11, 37, 65–70]. Unisensory responses are particularly sensitive to modality switches, often exhibiting RT costs following a switch that can rival the magnitude of the RSE itself. In contrast, multisensory responses tend to be less affected by the recent trial history (e.g., [65, 66]), leading to an asymmetry that can modulate the observed RSE and exaggerate RMI violations. As such, any comprehensive model of multisensory integration should consider these sequential effects.

In unisensory decision-making, recent models have begun to incorporate such dependencies. For instance, Caie et al. [59] introduced a gated accumulation model in which trial history influences the gating dynamics of sensory evidence accumulation. In future work, we propose extending the relay model in a similar direction by embedding history-dependent mechanisms into its initial stage. This extension would enable the model to capture trial history effects in unisensory conditions and generate testable predictions for multisensory responses without requiring additional parameter fitting.

Finally, the relay model provides a mechanistically transparent alternative to other race-based accounts of multisensory decision-making. For example, Otto and Mamassian [11] proposed a modified race model that introduces an unspecified source of noise between parallel processing streams. However, while the addition of noise permitted RMI violations, it is insufficient to reproduce the full shape of empirical RT distributions. To improve model fit, a correlation parameter captures the above mentioned trial-by-trial dependencies in unisensory RTs (for the link between the correlation parameter and the RMI, see [38]). Although both models can reproduce behavioural data, the relay model does so with computationally minimal and mechanistically explicit assumptions. By proposing a stage-based architecture governed by a single RT-share parameter, it offers a more interpretable and parsimonious account of multisensory integration dynamics.

In conclusion, the relay model provides a principled and computationally grounded framework for understanding the temporal dynamics of multisensory decision-making. By simply extending the classic race model to incorporate two sequential race stages, it accounts for the full magnitude of the RSE, including RMI violations, while remaining mechanistically transparent. Its modular design allows for integration with broader decision-theoretic models, including those that incorporate gating, evidence accumulation, and trial history effects. As such, the relay model provides a new tool for linking behavioural findings to underlying neural and cognitive processes in multisensory integration.

## Methods

### Data

The data presented in Figure 1b, along with the modelling results in Figures 2 and 3, are based on Experiment 1 from Miller [4]. In this experiment, 74 participants completed a redundant signals task involving three stimulus conditions: a visual stimulus (an asterisk at the centre of the screen), an auditory stimulus (a 780-Hz tone), or both stimuli simultaneously. Participants were instructed to press a key as quickly as possible upon detecting any stimulus. Each participant completed two blocks of 40 trials, with each block containing 10 trials per stimulus condition as well as 10 catch trials (no stimulus).

For analysis, RTs for each condition within each block were rank-ordered and then averaged across blocks and participants [4, 71]. This yielded three cumulative group RT distributions (auditory, visual, and audiovisual), each based on 1480 trials. As raw data are not available from the original publication, we extracted group-level data using graphical digitisation (Figure 1 in [4]). The figure was digitised, and the pixel-to-millisecond scaling was calibrated using a known 100 ms interval. RT values were then read directly from the scaled x-axis coordinates of the plotted points.

The data underlying Fig 4 are taken from Miller’s follow-up study [36]. Two participants completed the same redundant signals task as described above. Crucially, this study introduced temporal asynchronies between the auditory and visual stimuli. Audiovisual trials were divided into 11 conditions: a synchronous condition, five with the visual stimulus leading (SOAs from −33 to −167 ms), and five with the auditory stimulus leading (SOAs from 33 to 167 ms), all in 33 ms increments. Each participant completed 40 blocks of trials. Each block contained 170 trials, including 10 trials per stimulus conditions (13 conditions: auditory, visual, and 11 audio-visual SOAs) and 40 catch trials (no stimulus). All empirical data can be accessed from [72].

Summary statistics were reported for each condition and participant (Table 1 in [36]). These included the mean, standard error, and median of RTs, as well as the magnitude of the RSE. We used data only from participant B.D., whose responses showed significant RSEs in 7 of the 11 SOA conditions (based on 400 trials per condition). By contrast, participant K.Y.’s data showed significant RSEs in only three conditions, a finding that Miller described as “somewhat atypical in view of the consistency with which RSEs are obtained” [36].

### Unisensory RT distributions

Race models allow for parameter-free predictions of the RSE directly from the observed unisensory RT distributions. Basic predictions, such as those from Raab’s race model (probability summation, Eq 2) or the RMI (Eq 3), can be computed from empirical data without distributional assumptions (e.g., [73, 74]). However, generating more complex predictions, such as those required by the relay model, requires assuming a particular RT distributional form.

For all simulations, we modelled unisensory RTs using the IG distribution (also known as Wald distribution; [75]). The IG distribution is well-suited for modelling non-negative, positively skewed data such as RTs [32, 33]. Consequently, we assumed that unisensory RTs are IG-distributed with mean μ and shape parameter λ:

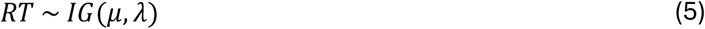

For the group-level RT distributions extracted from [4], we estimated the IG parameters for each unisensory condition separately: (μ_A_, λ_A_) for the auditory condition, and (μ_V_, λ_V_) for the visual condition. Parameters were optimised using MATLAB’s *fmincon* function, minimising the RMSE between the empirical and the simulated distributions at the 10 available quantiles.

For the SOA data extracted from [36], we estimated (μ_A_, λ_A_) and (μ_V_, λ_V_), using the simulated method of moments approach [76]. Here, the fitting procedure minimised the RMSE between the observed and simulated descriptive statistics (mean and standard deviation). Although [36] reports standard errors, we convert these to standard deviations (using n=400) to directly fit the standard deviation of the simulated distribution. The analytic mean was directly computed from μ and the standard deviation from 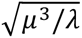.

The best-fitting parameters for both datasets are reported in Table 1. Importantly, all subsequent modelling work, whether for race model predictions or simulations under the relay model, was constrained by these parameters derived from the unisensory conditions.

### Stage-based representation of unisensory RTs

The relay model assumes that unisensory RTs are composed of two (or more) sequential processing stages (Fig 1c). Accordingly, the total RT can be expressed as:

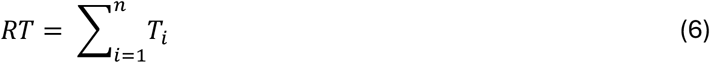

where *i* is the index of *n* stages.

We operationalised this assumption using the IG distribution, which was used to describe unisensory RTs as described above. A key mathematical property of the IG distribution is that the sum of IG-distributed random variables remains IG-distributed under specific scaling conditions. Formally, if:

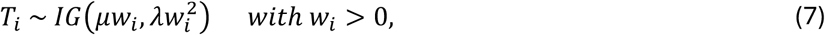

describes the processing time at sequential stages, then the total RT remains IG-distributed with mean μ and shape parameter λ (Eq 5). This property arises because the IG parameters are scaled with the weights *w_i_*, such that the convolution preserves the original distribution form. Thus, any unisensory RT distribution that is well-described by a single IG distribution can equivalently be expressed as the sum of multiple IG-distributed stages, with the mean and variance partitioned via the weights *w_i_*.

For model fitting and simulation purposes in the multisensory context, we imposed two constraints. First, we used normalised weights *w_i_* that sum up to one:

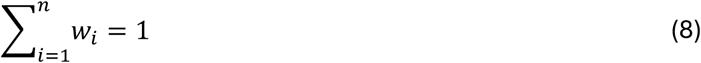

Thus, weights *w_i_* can be interpreted as the proportion of the total RT contributed by a specific stage (in terms of both mean and variance), which we describe as the RT share (Fig 2a). For example, *w*_1_ = 0.2 implies that the first stage accounts for 20% of the total RT. Stage 2 would then account for the remaining 80%.

Second, for simplicity, we assumed that the same RT share configuration applies to both audition and vision. For example, if the RT share at the first auditory stage is 20%, we assumed the same RT share for the corresponding visual stage. This simplification facilitates model fitting and interpretation but can be relaxed in future experiments designed to isolate modality-specific stage dynamics.

### Relay model

The basic relay model consists of two sequential race units (Fig 1c). For stage *i*, let the auditory and visual processing times be distributed as:

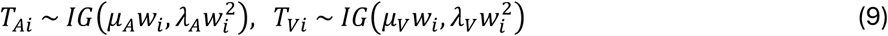

where *w_i_* represents the RT share of stage *i*. The corresponding CDFs are denoted *F_Ai_*(*t*) and *F_Vi_*(*t*), and their PDFs denoted *f_Ai_*(*t*) and *f_Vi_*(*t*), respectively. Because stage *i* completes as soon as either modality finishes processing, the duration of stage *i* in the multisensory case is:

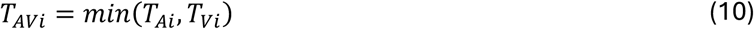

The CDF of *T_AVi_*, denoted *F_AVi_*(*t*), can be computed analogously to Raab’s race model under the assumption of statistical independence (Eq 2). The corresponding PDF, denoted *f_AVi_*(*t*), is given by:

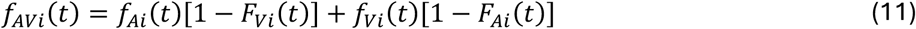

As the relay model assumes cross-modal initiation, the total multisensory *RT_AV_* is given by the sum of the durations of the sequential stages (Eq 4). To compute the CDF of *RT_AV_*, we convolved the CDF of first stage (*F_AV_*_1_) with the PDF of the second stage (*f_AV_*_2_):

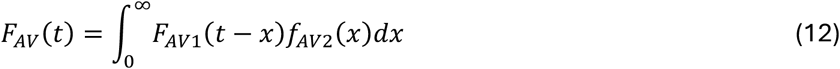

For simulations shown in Fig 3, we extended the relay model to more than two stages. The overall CDF for a multi-stage relay model was computed iteratively: starting with the CDF of the first stage, we successively convolved it with the PDFs of the additional stages. Each iterative computation is analogous to Eq 12.

For simulations shown in Fig 4, we applied the two-stage relay model to conditions involving SOA manipulations, as reported in [36]. To account for SOA manipulations, we introduced temporal lags into both stages. For example, when the auditory signal onset was delayed relatively to the visual signal (negative SOAs), we added a corresponding lag Δ*t* to the auditory process at the first stage:

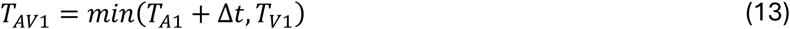

If the first stage completed before the auditory signal had even arrived, a residual delay was added to the auditory process at the second stage:

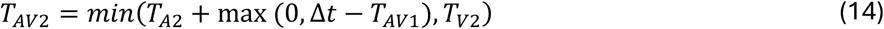

The model was adapted symmetrically for conditions in which the visual signal was delayed (positive SOAs). In both cases, the total multisensory RT is given by the sum of *T_AV_*_1_ and *T_AV_*_2_.

### RSE and RMI violations

To inspect the key features of the predictions derived from the relay model, we measured the RSE as well as violations of the RMI (Figs 2 and 3). To quantify these effects, we used geometric measures on the level of CDFs analogous to approaches described elsewhere [74, 77].

### Model fitting

In addition to running model simulations, we fitted the relay model to empirical data from [4, 36]. As outlined above, the relay model was constrained by the unisensory RT distributions estimated for each dataset (Table 1). To account for the observed multisensory RTs, the model included a single free parameter, the RT share of stage 1, which was constrained to the interval [0, 0.5].

To fit the relay model to the multisensory RTs from [4], we compared empirical quantiles of the multisensory condition to those generated from the relay model CDF (Eq 12). The relay model CDF was evaluated at the empirical quantile time points and the RMSE between the empirical and model CDF was minimised by adjusting the RT share parameter. Optimisation was conducted using MATLAB’s *fminbnd* function.

To fit the relay model to the multisensory RTs from [36], we used the relay model adapted for temporal asynchronies (Eqs 13 and 14). Miller [36] defined the RSE as the difference in mean RT between the faster unisensory and the multisensory condition. To generate model predictions, we sampled 100,000 RTs per condition and SOA and computed the RSE analogously. We minimised the RMSE between empirical and simulated RSEs across the 11 SOA conditions. Optimisation was conducted using MATLAB’s *bayesopt* function.

### Ethics Statement

The analysis carried out in this study involves model fitting of non-contentious anonymous secondary human data, and did not involve any new participant data collection (see Data). Ethical approval for data reuse was given by the University Teaching and Research Ethics Committee (UTREC) at the University of St Andrews (0740 - PS-0740-852-2025).

## Data Availability

All data and code used to produce the results and analyses presented in this manuscript is available on a GitHub repository at https://github.com/kalvinroberts/RelayModelCode. We have also used Zenodo to assign a DOI to the repository: <u>10.5281/zenodo.15830792</u>.

## Notes

### Competing Interest Statement

The authors have declared no competing interest.

